# Transfer of hepatocellular microRNA regulates cytochrome P450 2E1 in renal tubular cells

**DOI:** 10.1101/2020.01.06.895821

**Authors:** Olivia Matthews, Emma E Morrison, John D Tranter, Philip Starkey Lewis, Iqbal S Toor, Abhishek Srivastava, Rebecca Sargeant, Helen Rollison, Kylie P Matchett, Gillian A Gray, Chris Goldring, Kevin Park, Laura Denby, Neeraj Dhaun, Matthew A Bailey, Neil C Henderson, Dominic Williams, James W Dear

## Abstract

Extracellular microRNAs have been demonstrated to have the ability to enter kidney tubular cells and modify gene expression. We have used a Dicer-hepatocyte-specific microRNA conditional knock-out (Dicer-CKO) mouse to investigate functional microRNA transfer from liver to kidney under physiological conditions and in the context of drug toxicity. Dicer-CKO mice demonstrated a time-dependent decrease in the hepatocyte-derived microRNA, miR-122, in the kidney in the absence of other microRNA changes. During hepatotoxicity, miR-122 increased in kidney tubular cells; this was abolished in Dicer-CKO mice. Depletion of hepatocyte microRNAs increased expression and activity of the miR-122 target - cytochrome (CYP) P450 2E1 - in the kidney. Serum extracellular vesicles (ECVs) from mice with hepatotoxicity increased proximal tubular cell miR-122 and prevented cisplatin proximal tubular cell toxicity. miR-122 also increased in urinary ECVs during hepatotoxicity in humans. Transfer of microRNA was not restricted to liver injury – we detected miR-499 release with murine cardiac injury, and this correlated with an increase in the kidney. In summary, a physiological transfer of microRNA to the kidney exists, which is increased by liver injury. Regulation of renal drug response due to signalling by microRNA of hepatic origin represents a new paradigm for understanding and preventing nephrotoxicity.

## Introduction

MicroRNAs are key regulators of cellular gene expression.^1^ The primary microRNA transcript is cleaved in the cell nucleus to release a pre-microRNA that is exported into the cytoplasm. The cytoplasmic enzyme Dicer cleaves the pre-microRNA to produce a microRNA duplex, one strand of which is loaded onto an Argonaute protein to form the RNA silencing complex.^1^ In the circulation, cell-free microRNAs are protected from degradation by extra-cellular vesicles (ECVs) and protein complexes such as Argonaute,^2^ and they are potential disease biomarkers.^3^ Extra-cellular microRNAs can enter cells and change gene expression and cellular function *in vitro*.^4^ There is evidence that transfer of microRNA between organs occurs *in vivo* in pre-clinical models of diseases such as cancer.^5^ Therefore, extra-cellular microRNAs may represent a new class of signalling molecules. In this paper, we tested the hypothesis that microRNAs produced in the liver are transferred to regulate gene expression in the kidney. We focussed on the kidney because our earlier work demonstrated that microRNA species can enter kidney tubular cells under physiological hormonal control and modulate their mRNA targets.^6^ Also, there is a clinically important relationship between acute liver injury and kidney injury. For example, kidney function forms an integral part of all the clinical risk stratification models that are used to make decisions regarding need for liver transplantation to prevent death in patients with acute liver failure.^7^

MicroRNA-122 (miR-122-5p, miR-122) is highly expressed in hepatocytes (∼40,000 copies per cell – around 70% of total hepatocyte microRNA). ^8 9^ In patients with acute liver injury, and in pre-clinical models, the circulating concentration of miR-122 is increased 100-1000 fold.^10^ In over 1000 patients we have demonstrated that miR-122 is a sensitive and specific biomarker of acute liver injury risk after paracetamol (acetaminophen) overdose, the most common cause of acute liver failure in the Western world.^11^

The cytochrome P450 (CYP) enzymes, particularly CYP2E1, are responsible for generating the toxic metabolite (*N*-acetyl-*p*-benzoquinone imine - NAPQI) that causes cell death in the context of paracetamol toxicity.^12^ In the liver, CYP2E1 is established as being regulated by miR-122.^13, 14^ In the kidney, CYP2E1 is expressed in tubular cells, a cellular location where it can also mediate drug toxicity. For example, although rare, paracetamol is well recognised to directly produce acute kidney injury in the absence of liver injury.^15^ CYP2E1 deletion in mice prevents cisplatin-induced acute kidney injury demonstrating a central role for CYP2E1 in this established model of nephrotoxicity.^16^ In this paper we explore whether liver to kidney microRNA transfer regulates kidney CYP2E1 expression and activity, and whether liver-derived microRNA can modulate nephrotoxic tubular cell injury.

## Results

### Basal miR-122 in the kidney originates from the liver

To determine whether liver hepatocellular microRNA is transferred to other organs, we depleted hepatocyte microRNA by treating Dicer^flox/flox^ mice with a Cre recombinase-expressing adenovirus (AAV8-Cre) (or control AAV8 which did not express Cre (AAV8-null)). Exposure to AAV8-Cre resulted in a time-dependent decrease in liver Dicer expression at the mRNA and protein level (Figure S1). AAV8-null had no effect on Dicer. Liver depletion of Dicer resulted in a time-dependent decrease in miR-122, miR-192 and miR-151 (microRNA species enriched in the liver^17^) (Figure 1). There was no significant microRNA change following AAV8-null injection. To confirm miR-122 depletion we performed in situ hybridisation (ISH) on liver tissue, which demonstrated reduced tissue expression with AAV8-Cre treatment (Figure S2). In serum, miR-122 concentration was reduced when AAV-Cre treated mice were compared with AAV8-null (normalised to miR-39. Week 3 after AAV8 injection: AAV8-null median 7.4 (IQR 3.3-54.7); AAV8-Cre 1.3 (0.2-3.1). Week 4 after AAV8 injection: AAV8-null 4.8 (IQR 0.3-88.1); AAV8-Cre 0.3 (0.1-1.5) N=5 per group).

**Figure 1.**
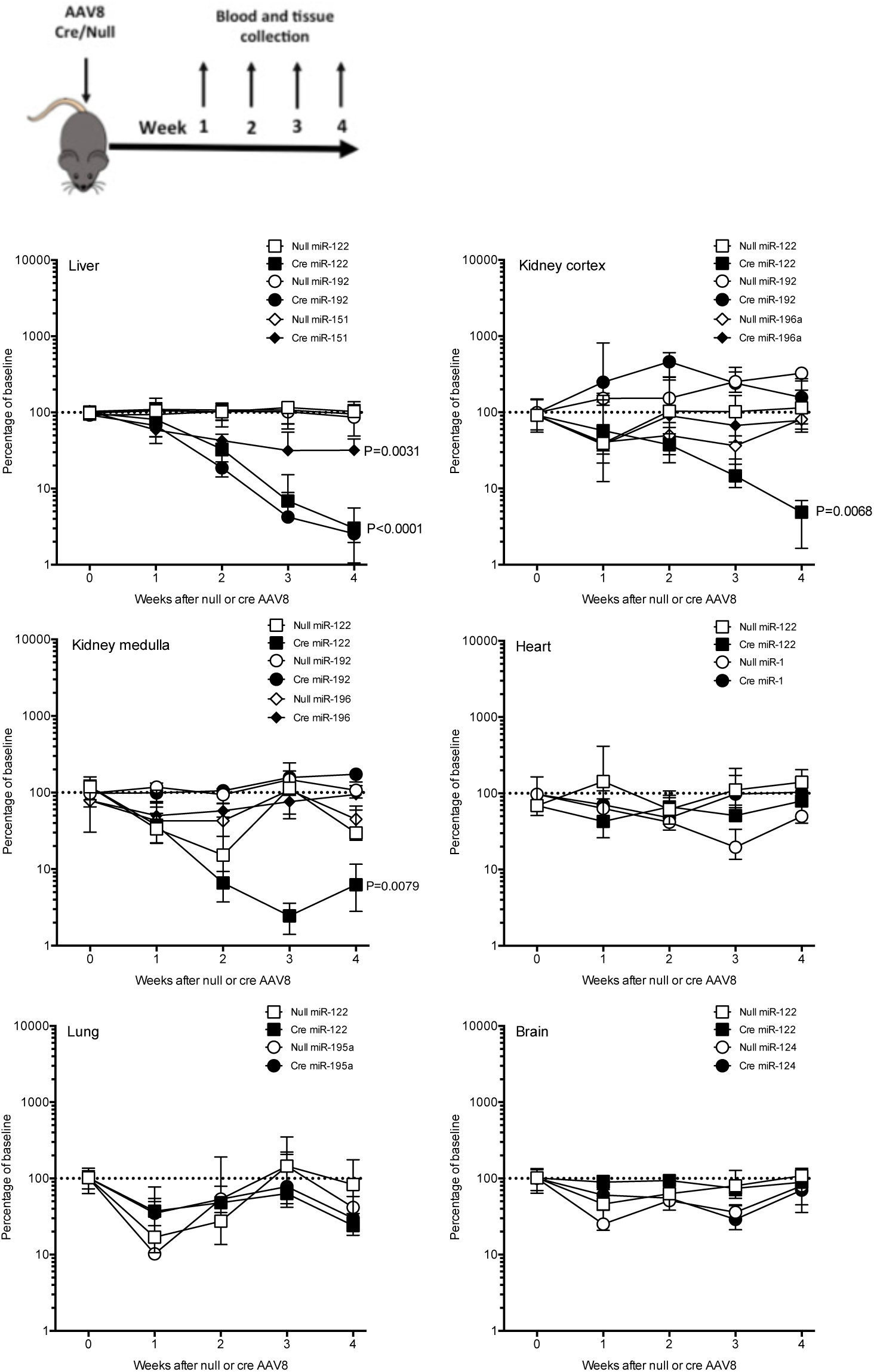
The percentage changes over time in organ microRNA expression after treatment of Dicer^flox/flox^ mice with AAV8 vector expressing or not expressing Cre Recombinase (Cre). Schematic for study design is presented. In graphs, for each organ, the expression of microRNAs is expressed as a percentage of the untreated Dicer^flox/flox^ mice (baseline). Data are normalised to U6. Filled symbols represent mice receiving AAV8-Cre. Unfilled symbols represent AAV8-null. Symbols represent the time point median and the error bars define the inter-quartile range. N=20 for each AVV8/microRNA combination (N=5 for each time point). Linear regression was performed then for each microRNA Cre treatment was compared to null treatment to determine whether slopes were significantly different.

**Figure 2.**
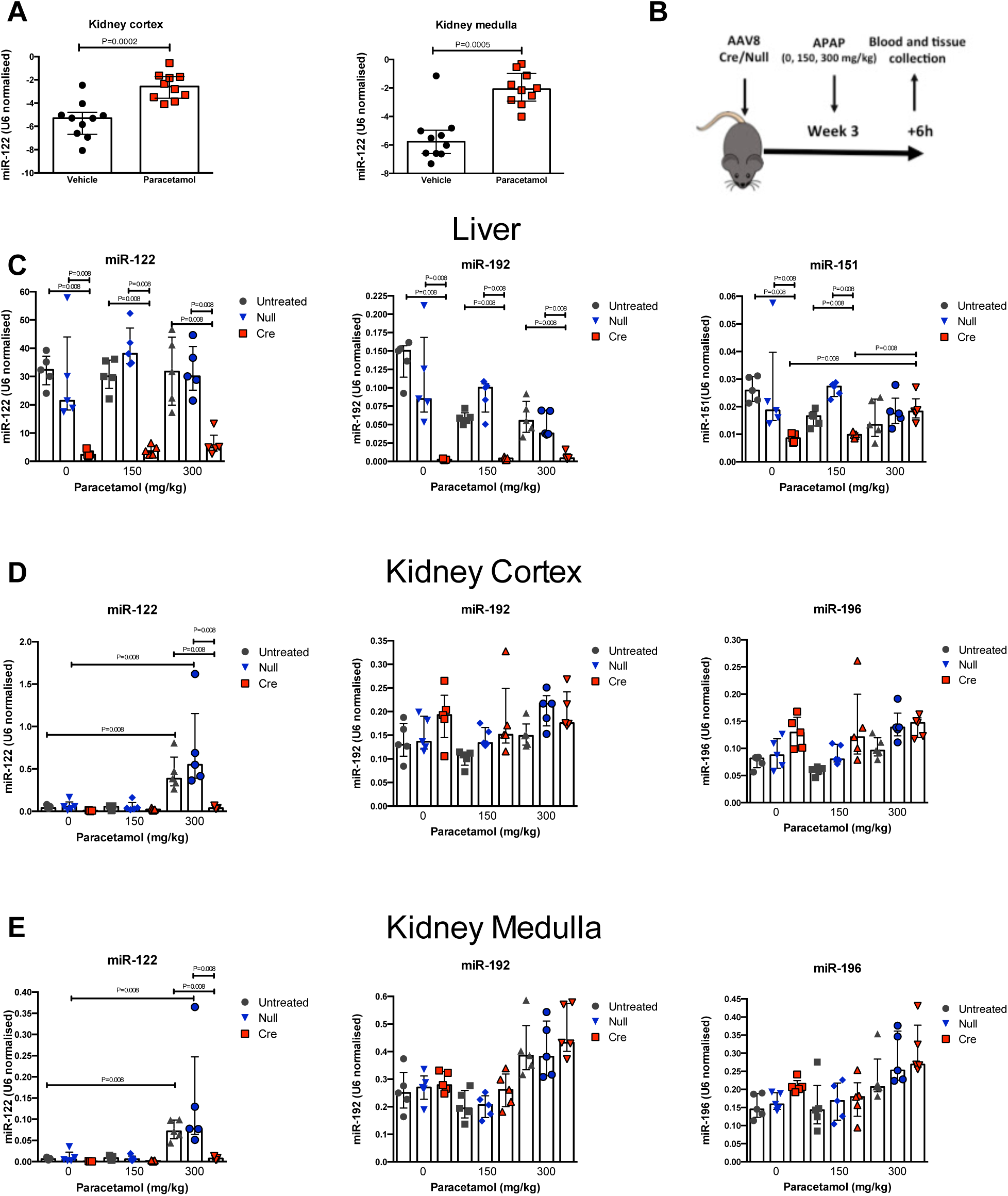
A) miR-122 was increased in the kidney cortex and medulla 6 hours after mice were treated with paracetamol (300mg/kg i.p.). N=10 per group. B) Schematic overview of study (APAP = paracetamol). C-E) Dicer^flox/flox^ mice were treated with AAV8 vector expressing or not expressing Cre recombinase (Cre or null). 3 weeks after AAV8 treatment mice received paracetamol 150 or 300 mg/kg (or vehicle (0)) ip, then liver (C), kidney cortex (D) and kidney medulla (E) were harvested 6 hours later. Untreated = Dicer^flox/flox^ mice not receiving AAV8. MicroRNA expression is expressed as U6(ct) - miR (ct). N=5 per group. Statistical significance was determined by Mann-Whitney Test. Data are represented individual mice with bars representing median and IQR.

The liver specificity of Cre delivery by the AAV8 vector was demonstrated by there being no change in Dicer expression in the kidney (cortex and medulla), heart or lung (Figure S1). There was a decrease in brain Dicer following AAV8-Cre. miR-122 concentration in these non-hepatic organs was determined alongside selected, organ-enriched, microRNAs (kidney: miR-192, miR-196a; heart: miR-1; lung: miR-195a; brain: miR-124).^9^ The only microRNA to significantly decrease following AAV8-Cre treatment of mice was miR-122 in the kidney cortex and medulla (Figure 1). To determine whether miR-122 was transcribed in the kidney we measured the primary miR-122 transcript. It was barely detectable in the kidney cortex and medulla (expression normalised to *Gapdh* – kidney cortex median 0.0023 (IQR 0.0016-0.003), in contrast to liver pri-miR-122 1.7 (0.9-5.4) N=5 per group) and was unaffected by AAV8 treatment (3 weeks after AAV8: null 0.0035 (0.0028-0.0044). Cre 0.0037 (0.0026-0.005) N=5 per group). Therefore, the change in miR-122 in the kidney was not due to a change in *de novo* synthesis.

### Increased miR-122 transfer from to the kidney occurs following liver injury in mice and humans

We treated wild-type C57BL/6 mice with an hepatotoxic dose of paracetamol (300mg/kg). Published studies demonstrate substantially increased circulating miR-122 6 hours after dosing, so this time window was chosen to study transfer to the kidney.^18^ In our study, after 6 hours, paracetamol resulted in centri-lobular hepatocyte necrosis and an elevation in serum ALT activity (300mg/kg dose, around 38 fold) and serum miR-122 concentration (around 18-fold) (Figure S3). Interestingly, the liver tissue expression of miR-122 was significantly altered after paracetamol exposure. There was loss of miR-122 from necrotic centri-lobular hepatocytes but a marked increase in expression in the areas surrounding the necrotic cells with the highest expression in the viable hepatocytes closest to the injury (Figure S2). Consistent with hepato-renal transfer, the concentration of miR-122 increased in the kidney cortex and medulla (Figure 2). To confirm that the liver was the source of the increase in miR-122 in the kidney we repeated the experiment with Dicer deletion using the AAV8Cre/Null (or no AAV8) followed 3 weeks later by paracetamol exposure (0, 150 and 300 mg/kg). In this model paracetamol treatment resulted in histological and biochemical liver injury with higher ALT activity and increased necrosis in the AAV8-Cre treated group. After AAV8-Cre treatment, miR-122 expression in the liver tissue following paracetamol exposure was lost in the areas surrounding necrotic cells (Figure S2). As in the wild-type C57BL/6 mice, there was a significant increase in kidney cortex and medulla miR-122 in untreated and AAV8-null treated Dicer^flox/flox^ mice. This increase was substantially attenuated when the Dicer^flox/flox^ mice were treated with the AAV8-Cre vector (14-fold difference when AAV8-Cre compared to AAV8-null after 300mg/kg paracetamol) (Figure 2). There was no effect on Dicer in the kidney (Figure S4) and no change in miR-192 or miR-196a (Figure 2). There was no expression of the primary miR-122 transcript in the kidney cortex and medulla without or with paracetamol treatment in any treatment group (Ct value >35). To determine which kidney cell type had increased miR-122 after liver injury, we FACS sorted kidney cells. There was a significant increase only in the miR-122 content of LTL+ tubular cells (Figure S5).

**Figure 3.**
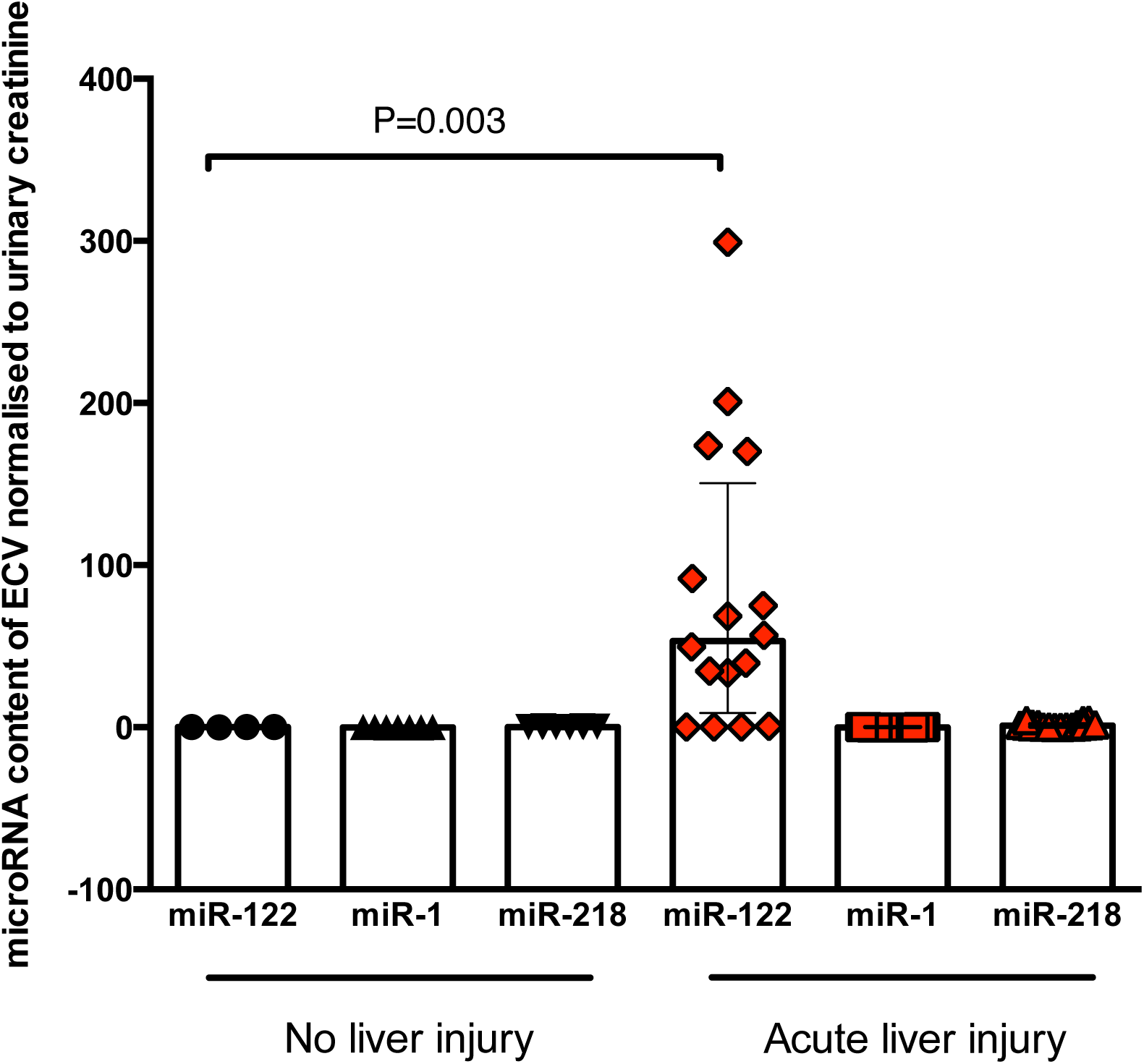
Urinary extra-cellular vesicles (ECVs) contain miR-122, which was significantly increased in patients with paracetamol-induced acute liver injury. Urine was collected from patients who had taken a paracetamol overdose that required treatment but did not cause liver injury (’No liver injury’) and patients with liver injury (ALT>1000U/L). ECVs were isolated by ultra-centrifugation. MicroRNA concentration was measured and expression normalised by urinary creatinine. Statistical significance was determined by Mann-Whitney Test. Data are represented individual mice with bars representing median and IQR.

**Figure 4.**
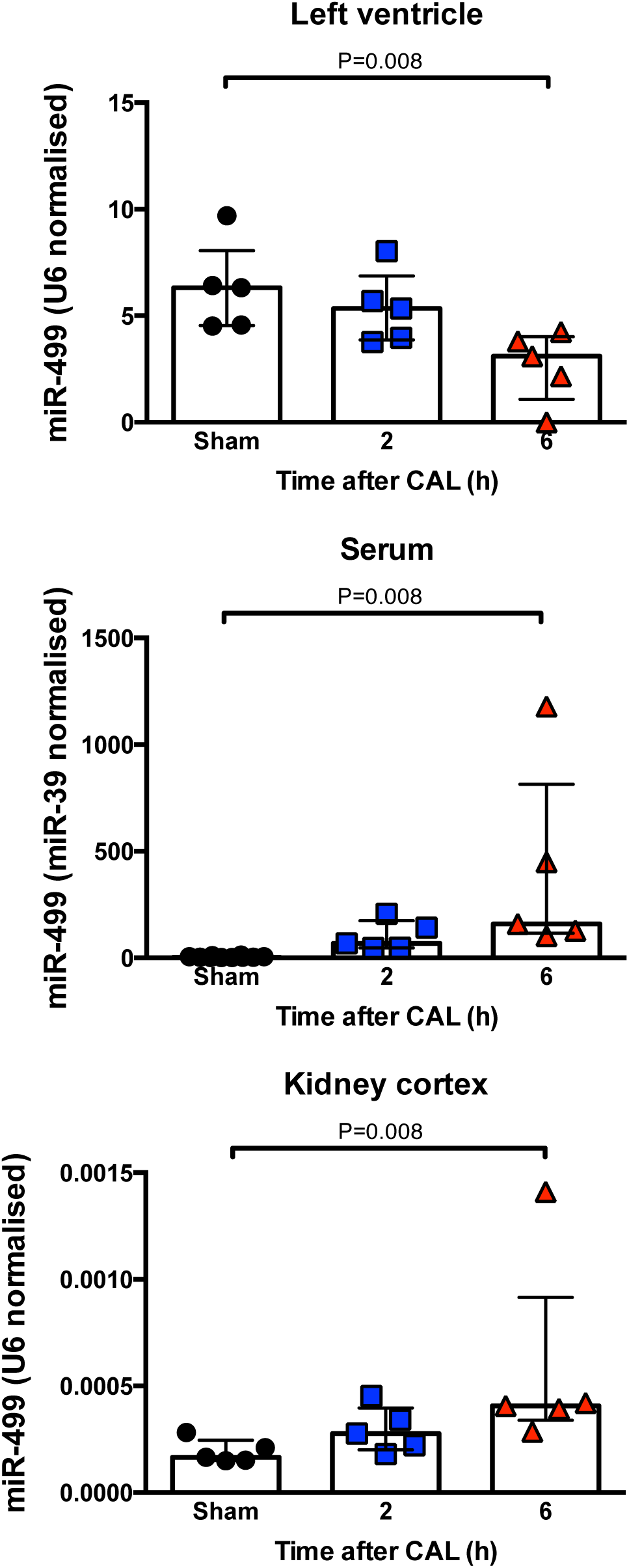
Myocardial injury was induced in mice by coronary artery ligation (CAL). After 2 or 6 hours tissue and serum was collected. Sham operated mice had thoracotomy performed but no ligation. In the left ventricle and kidney cortex miR-499 expression is expressed as U6(ct) - miR (ct). In serum miR-499 is normalised by spike in miR-39. Statistical significance was determined by Mann-Whitney Test. Data are represented individual mice with bars representing median and IQR.

**Figure 5.**
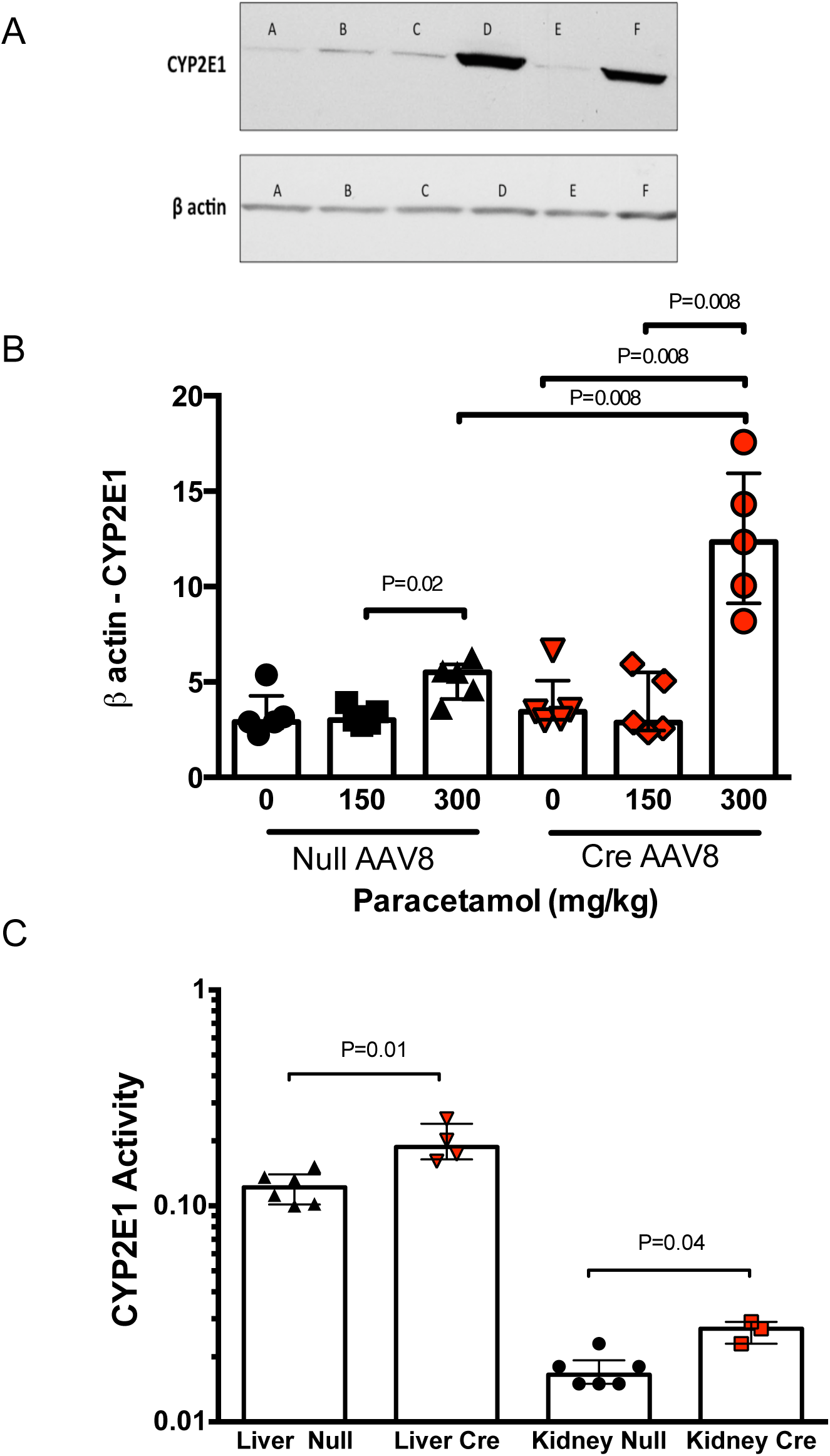
Dicer^flox/flox^ mice were treated with AAV8 vector expressing or not expressing Cre recombinase (Cre or null). A) Western blot of kidney cortex CYP2E1 A, C, E. mice treated with AAV8 null then kidney collected 1, 2 or 3 weeks later. B, D, F mice treated with AAV8 Cre then kidney collected 1, 2 or 3 weeks later B) Cytochrome P450 2E1 (CYP2E1) mRNA expression is expressed as b-actin(ct) - CYP2E1 (ct). 3 weeks after AAV8 treatment mice received paracetamol 150 or 300 mg/kg (or vehicle (0)) ip, then the kidney cortex was harvested 6 hours later. C). CYP2E1 activity was determined by metabolism of chlorzoxazone. Statistical significance was determined by Mann-Whitney Test. Data are represented individual mice with bars representing median and IQR.

To begin to translate this mouse work into humans, we measured miR-122 in urinary ECVs from patients following paracetamol overdose (without and with acute liver injury, serum ALT activity >1000U/L). miR-122 was substantially increased in the urinary ECVs of those patients with liver injury demonstrating transfer of miR-122 from circulation to the urinary space in humans with paracetamol hepatotoxicity (Figure 3). There was no change in two microRNAs enriched in non-hepatic organs.

### Cardiac microRNA is transferred to the kidney following myocardial ischaemia

To determine whether microRNA transfer to the kidney is restricted to liver injury we explored kidney microRNA expression in a model of ischaemic cardiac injury induced by coronary artery ligation (CAL). Cardiac injury was demonstrated by an increased circulating troponin concentration 6 hours after CAL (Sham: 1.1ng/mL (0.9-1.9). CAL: 22.4ng/mL (17.0-24.5) P=0.016 N=5). A panel of microRNAs were measured in the circulation and miR-499 was identified as the species with the largest fold increase with CAL (relative fold increase from baseline 159 (116-813), data not shown for other species). This increase in the circulation was associated with a significant decrease in the expression of miR-499 in the left ventricle of the heart and a concurrent increase in the kidney (Figure 4). This demonstrates that microRNA released from the heart could also be transferred to the kidney.

### Kidney CYP2E1 is regulated by liver-derived microRNA

CYP2E1 expression was determined by immunohistochemistry (IHC). Increased expression was observed in the liver treated with AAV8-Cre compared with AAV8-null (Figure S6), which is consistent with published papers demonstrating that the cellular expression of CYP2E1 is regulated by miR-22.^13, 14^ To confirm that enzyme activity is regulated by liver microRNA we used the chlorzoxazone probe to assay CYP2E1 activity in liver-derived microsomes. There was a significant increase in liver enzyme activity in Dicer ^flox/flox^ mice treated with AAV8-Cre compared to AAV8-null treatment (Figure 5C).

**Figure 6.**
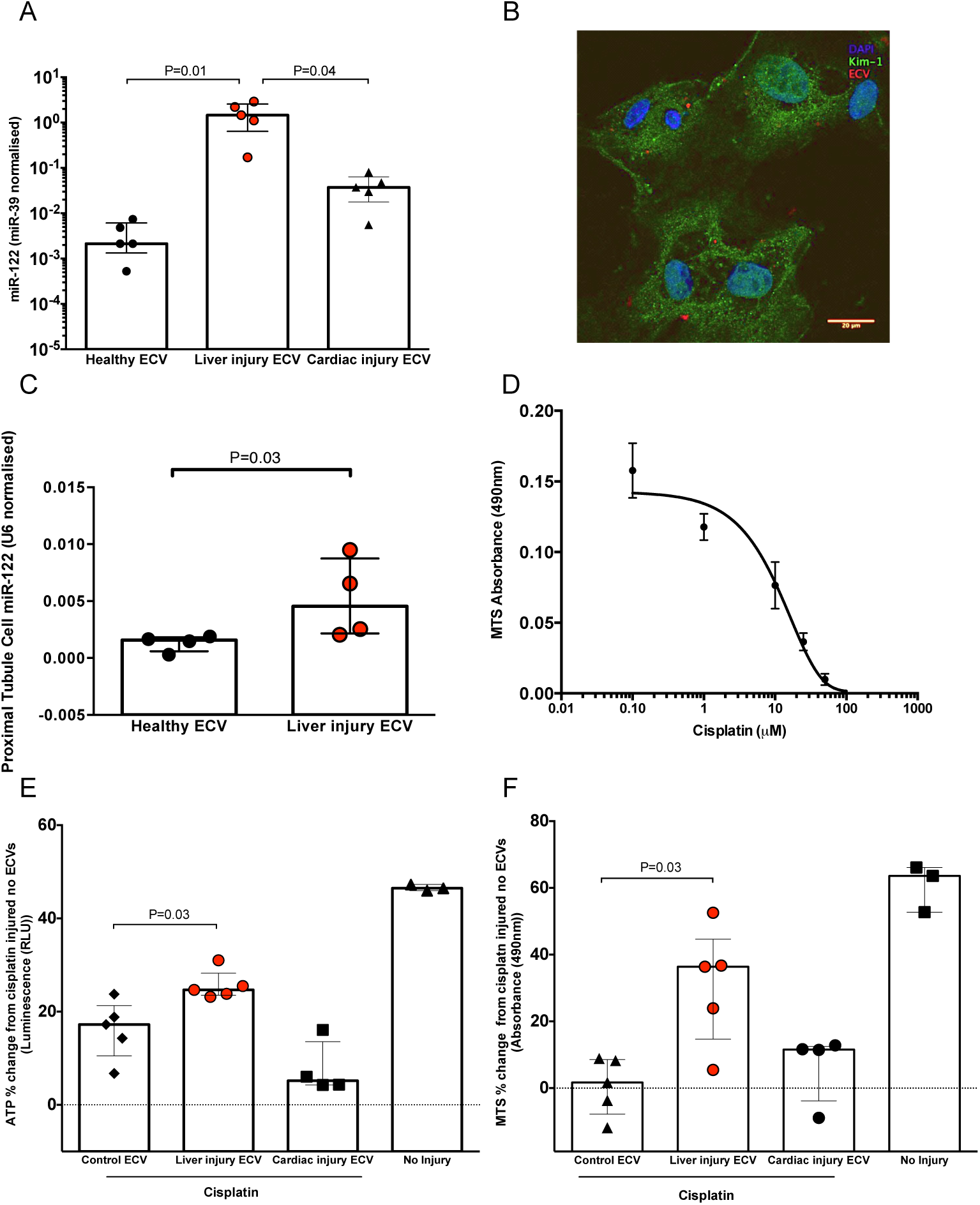
A) miR-122 in ECVs from serum of healthy mice, mice with liver injury (300mg/kg paracetamol IP) or cardiac injury (induced by CAL). B). Murine primary proximal tubular (PTT) cells internalize ECVs (red). Green represents KIM-1, an archetypal proximal tubular protein. Blue stain is DAPI. C) ECVs were isolated from the serum of mice treated with a toxic dose of paracetamol (300mg/kg ip - liver injury ECV) or healthy mice. Equal ECV numbers were applied to PTT cells and the miR-122 content of the cells was determined. D) PPT cells were exposed to cisplatin for 48 hours prior to assessment of cell viability by NADPH activity (N=4). E & F). PTT cells were co-incubated with circulatory ECVs (250 x10^8^/ml) for 48 hours and subsequently injured with 10 μM cisplatin. Circulatory ECVs were derived from healthy mice (control), liver injury mice (300mg/kg paracetamol IP) or cardiac injury induced by CAL. Cells that were not injured with cisplatin and had no additional ECVs added are labeled no injury. All results were normalised to the results from PTT cells injured with 10 μM cisplatin with no additional ECVs (baseline). Data represents ATP concentration (Figure E) and NADPH activity (Figure F) as percentage change from baseline. Statistical significance was determined by Mann-Whitney Test. Data are represented individual mice with bars representing median and IQR.

In the kidney tissue, CYP2E1 was localised to tubular cells in the cortex (Figure S6). In Dicer ^flox/flox^ mice pre-treated with AAV8-Cre, CYP2E1 was significantly increased at the RNA and protein level compared to AAV8-null in untreated mice and after paracetamol exposure (Figure 5 A&B). Furthermore, there was increased CYP2E1 activity in kidney-derived microsomes from AAV8-Cre treated mice (Figure 5C).

ECVs containing miR-122 were isolated from mouse serum. The miR-122 concentration of the ECVs from mice with liver injury due to paracetamol was around 460 and 39 fold higher than untreated and cardiac injury ECVs, respectively (Figure 6A). Fluorescently-labelled ECVs from liver injury mice entered mouse primary proximal tubular cells (Figure 6B) and resulted in a significant increase in proximal tubular cell miR-122 concentration (Figure 6C). To determine whether microRNA release from the liver could prevent drug toxicity in the kidney we cultured mouse primary proximal tubular cells with equal numbers of ECVs isolated from the following groups: circulation of untreated mice, mice with liver injury induced by paracetamol or mice with cardiac injury induced by CAL. ECV treatment for 48 hours was followed by exposure of the cells to cisplatin (the toxicity of which is dependent on CYP2E1 activity^16^) at the EC50 toxicity dose (Figure 6D). The ECVs from mice with liver injury significantly attenuated cisplatin toxicity. By contrast, ECVs from healthy mice and mice with cardiac injury had no effect (Figure 6E&F).

## Discussion

The work described in this paper demonstrates that miR-122 is transferred from the liver to the kidney tubular cells as a physiological process (in the uninjured mouse) that is increased by acute liver injury. MicroRNA is also transferred from the injured heart to the kidney. In the kidney, microRNA originating from the liver regulated CYP2E1 RNA and protein expression and activity and attenuated cisplatin toxicity. In humans with paracetamol toxicity there is evidence of increased miR-122 in urinary ECVs, which supports our mouse data being a faithful reflection of human pathophysiology.

miR-122 has two unique properties which facilitated our experiments. Firstly, it is highly enriched in hepatocytes compared to other organs. Data from the FANTOM consortium demonstrates that miR-122 is approximately 18,000 higher in hepatocytes than renal proximal tubular cells.^9^ Secondly, it is released in large amounts from the injured liver into the circulation. These properties allowed us to use miR-122 to track endogenous microRNA transfer by liver-specific deletion of the enzyme that generates mature microRNA species (Dicer). This approach also allowed us to confidently exclude off-target Cre recombinase activity in non-hepatic organs by measurement of Dicer and organ-selective microRNAs. When Dicer was deleted in the liver there was a subsequent substantial miR-122 decrease in the kidney. The primary transcript for miR-122 was undetectable in the kidney. The FANTOM consortium used RNA-sequencing and cap analysis of gene expression (CAGE) across multiple primary cell types. This demonstrated the presence of mature miR-122 in kidney proximal tubular cells without expression of its promoter.^9^ This supports the conclusion from our studies - miR-122 expression in the kidney results from transfer from the liver. This is consistent with the results of Rivkin *et al.* who used macrophage depletion or antagomirs to indirectly inhibit LPS-induced miR-122 release from mouse hepatocytes and reported a corresponding reduction of kidney miR-122.^19^ The data presented in our paper now demonstrate that microRNA transfer from the liver to the kidney occurs as part of normal physiology and does not require experimental induction of inflammation or tissue injury. It is likely that multiple microRNA species are transferred from liver to kidney as part of normal physiology. The unique properties of miR-122 described above allow us to detect a change whereas other microRNAs will be synthesised locally in the kidney. This local expression is likely to mask any change due to decreased transfer from the liver (a hypothesis supported by miR-192 not changing in the kidney despite a decrease in the liver). Future studies should define the total contribution of liver-derived RNA in the kidney.

Circulating microRNAs change with tissue injury as exemplified by the increase in miR-122 that accompanies liver injury. In this paper we demonstrate that miR-122 released as a result of paracetamol toxicity targets the kidney tubular cells. Similar transfer to the kidney was demonstrated using a model of ischemic injury to the myocardium with miR-499 as the target microRNA. This demonstrates that the kidney is a key node that internalises microRNA released from injured organs. Recent *in vivo* data demonstrate that miR-122 released from the injured liver can induce acute lung inflammation by activating alveolar macrophages.^20^ In our studies there was not a significant decrease in lung miR-122 following liver microRNA depletion. In contrast to the kidney, this suggests that microRNA transfer from the liver to the lung is not significant in the healthy uninjured state. In humans, the cargo of miR-122 in urinary ECVs is substantially increased when the liver is injured which supports our mouse model of liver to kidney transfer occurring in human disease. The mechanism by which miR-122 goes from the circulation to urinary ECVs remains to be defined. It could be that ECVs transfer from blood to urine or miR-122 containing ECVs are generated *de novo* by the kidney tubular cells when the liver is injured.

There are many different pathways that could be regulated in the kidney by microRNA originating from other organs and this will be a subject of further research. We focussed on CYP2E1 because it is established as being regulated by miR-122 and it is a key enzyme in the drug metabolism that underlies liver and kidney toxicity. Our data demonstrate that CYP2E1 RNA and protein expression and enzymatic activity is regulated by liver-derived microRNA in the liver and the kidney. As others have reported, the consequence of microRNA depletion in the liver was increased injury following paracetamol exposure. ^13, 14^ Interestingly, in the liver, miR-122 was substantially increased in the hepatocytes surrounding the necrotic core (miR-122 expression was lost from the core). This effect was not apparent when bulk tissue expression was measured by PCR. Theoretically, transfer of miR-122 may reduce drug toxicity in the kidney by down-regulating CYP enzyme activity. Consistent with this, ECVs from mice with liver injury prevented cisplatin toxicity, which is dependent on CYP2E1 activity. John *et al.* reported that patients with acute liver failure who die have lower circulating miR-122, which is consistent with miR-122 inter-organ transfer having a possible protective effect in humans.^21^

In summary, there is physiological transfer of microRNA from the liver to the kidneys that is increased by hepatotoxicity and regulates kidney CYP enzymes. miR-122 represents the majority of the hepatocyte microRNA cargo and, following hepatotoxicity in humans, it is the highest concentration microRNA in the circulation. We propose miR-122 release (and potentially other microRNA species) has the potential to mediate resistance to drug toxicity in the kidney and may be, therefore, acting as a break on multi-organ failure in the context of drug-induced liver injury.

## Methods

### Animal Studies

Mice with free access to standard chow and water were housed in groups of 4-8 in open-top cages, at 22°C ± 1°C, 55% humidity and on a 12 hour light dark cycle (lights on at 07:00). The mice were allowed to acclimatise to the environment for at least a week prior the start of each experiment. At the end of the study, mice were euthanized by rising CO_2_ exposure followed by exsanguination, unless stated otherwise.

#### Dicer conditional knock-out (Dicer-CKO)

Dicer^flox/flox^ mice (male and female, 2-3 months old) (B6.Cg-*Dicer1^tm1Bdh^*/J strain) were injected with a single tail vein injection of the hepatocyte-specific AAV8.TBG.PI.Cre.rBG (AAV8-Cre) or AAV8.TBG.PI.Null.bGH (AAV8-null) (Penn Vector Core, Pennsylvania, USA, 2.5×10^11^- 6.25×10^10^ viral genomes/100µl dose in sterile PBS). Untreated mice Dicer^flox/flox^ (baseline) were used as controls where indicated. After 1-4 weeks, blood and tissue were collected. Whole blood was spun for 10 minutes at 8000 x *g*, 4°C, serum was isolated and stored at −80°C. Tissue was placed in 4% paraformaldehyde (Sigma Aldrich) for 24 hours, followed by storage in 70% ethanol and finally paraffin embedded for sectioning. The rest of the tissue was put into RNAlater (Sigma Aldrich) and snap frozen for long-term storage at −80°C. Prior to tissue collection the circulation was perfused with sterile saline. Where indicated in the results section, mice were treated with paracetamol as described below.

#### Paracetamol hepatotoxicity model

Wild-type male C57BL/6JCrl mice (2-3 months old) or Dicer-CKO mice were fasted overnight for 12 hours then intraperitoneally injected with paracetamol (ACROS organics, Geel, Belgium) dissolved in sterile PBS (Sigma Aldrich, Dorset, UK). Tissue and blood collection occurred 6 hours after paracetamol injection. Prior to tissue collection the circulation was perfused with sterile saline. In specific experiments, circulatory ECVs were isolated from mouse serum as described previously ^22^. In brief, mouse serum was vigorously vortexed then centrifuged at 12,500 x *g* for 30 minutes. The supernatant was then centrifuged at 120,000 x *g* for 70 minutes to pellet the ECV fraction. The pellet was washed and then re-centrifuged before final resuspension in PBS. In certain studies pelleted ECVs were conjugated with Cell Tracker 655 (Invitrogen, CA, USA) following the manufacturer’s protocol. ECV size distribution and number was measured by nanoparticle tracking analysis as previously described.^23^ Mouse plasma alanine transaminase activity (ALT) was determined using a commercial serum ALT kit (Alpha Laboratories Ltd., Eastleigh, UK) adapted for use on either a Cobas Fara or Cobas Mira analyser (Roche Diagnostics Ltd, Welwyn Garden City, UK). In certain studies, kidney cells were isolated by fluorescence-activated cell sorting (FACS). Kidneys were dissociated in RPMI media containing collagenase II (Sigma-Aldrich C6885), collagenase D (Roche 11088858001), dispase (Gibco 17105041) and DNase I (Roche 04716728001). Cells were then incubated with BD Fc Block™ purified anti-mouse CD16/CD32 (BD Biosciences 553141) prior to incubation with fluorescein-labelled *lotus tetragonolobus lectin* (LTL, Vector Laboratories FL-1321), Brilliant Violet 605™ anti-mouse CD31 (Biolegend 102427) and APC rat anti-mouse CD45 (BD Bioscience 561018). FACS was performed on the BD FACSAria™ II apparatus (BD Biosciences).

#### Coronary artery ligation (CAL) model

CAL was performed in wild-type C57/BL6 mice as described previously ^24, 25^. In brief, an incision was made in the lower thorax and the chest opened at the fourth intercostal space and held open using retractors. The left anterior descending coronary artery was ligated in mice randomised to the CAL group, at the level below the left atrium. After ligation, the chest was closed. For mice undergoing sham surgery, thoracotomy alone was performed; ligation of the left anterior descending coronary artery was not conducted. An ultra-sensitive mouse cardiac troponin-I ELISA (Life Diagnostics, Stoke on Trent, UK) was used on serum as per the manufacturer’s instructions.

### Human studies

Adult patients (age>16 years) admitted to the Royal Infirmary of Edinburgh, UK following paracetamol overdose without or with acute liver injury (ALT>1000U/L) were included. Urine was collected and ECVs were isolated from the whole urine samples as previously described.^23^

### In vitro studies

#### Primary proximal tubular cell isolation

Male mice were euthanised and kidneys were removed immediately under aseptic conditions and the cortex macroscopically dissected. The cortex was minced and incubated with 1mg/ml collagenase I and IV (Sigma Aldrich, Dorset, UK) for 45 minutes at 37 °C. The resultant solution was ground and serially sieved to a final filter size of 40µm. The filtered solution was centrifuged at 27 000 x *g* through a 48% Percoll gradient (Sigma Aldrich, Dorset, UK) and each of the 4 distinct bands were carefully removed. Cells contained within the lowermost band (F4) were washed and passed through a 40µm sieve. The cells were resuspended in DMEM/Hams’s F12 media with glutamax containing: 5 μg/ml insulin, 50 nM hydrocortisone, 10 ng/ml EGF, 5 μg/ml transferrin, 50 nM sodium selenite, 10 nM triiodothyronine, 100 U/ml penicillin, 100 μg/ml streptomycin and 1% (wt/vol) exosome depleted FBS (System Biosciences, CA, US). Studies of NHE3 and KIM-1 protein expression and alkaline phosphatase activity confirmed the presence of proximal tubular cells (PPT) in the F4 fraction (data not shown). PPT cells were seeded on at a cell density of 5×10^3^ cells/well for 48 hours. Then circulatory ECVs were co-incubated with cells at a concentration of 250 x10^8^/ml media, 48 hours prior to induction of injury. Nephrotoxic injury was induced by the addition of cisplatin (Cambridge Bioscience) to PPT cells for 48 hours. Experiments were conducted in technical triplicate (3 wells) and biological quintuplicate (ECVs isolated from 5 animals per group). Cell viability was determined by the CellTiter-Glo^®^ cell viability assay (ATP quantification) (Promega, Wisconsin, USA) assay and the CellTiter 96^®^ Aqueous Assay (MTS assay) (Promega, Wisconsin, USA) as per manufacturer’s instructions.

#### RNA preparation and qRT-PCR

Total RNA was extracted and purified from serum using the miRNeasy Serum/Plasma Kit (Qiagen) with *C.elegans* miR-39 (Qiagen) added as an external control. Tissue total RNA (250ng) was extracted and purified using the miRNeasy Mini Kit (Qiagen). cDNA was synthesised using the miScript II RT kit (Qiagen) according to the manufacturer instructions and diluted 1:10 to perform qRT-PCR. MicroRNA and mRNA quantification was carried out using the miScript SYBR green PCR kit (Qiagen) on the Lightcycler 480 (Roche Diagnosis). Quantitect and miScript primers were purchased from Qiagen. For the quantification of the primary miR-122 transcript, the Taqman Gene Expression Mix was used for the detection of primary miR-122 transcripts (Life Technologies).

#### Histological scoring (necrosis quantification)

Liver H&E sections were scored for injury as follows: 100x (distance (µm) between the central vein and edge of the necrotic zone)/ (total distance (µm) between the central vein and portal triad). 10 measurements chosen at random were made per section.

#### Immunohistochemistry

CYP2E1 immunohistochemistry in the liver and kidney was performed on the Discovery ULTRA Staining Module (Roche Diagnostics). For staining preparation, the sections were exposed to 100°C (4 mins) and incubated with Ventana Cell Conditioner 1 (5x 8mins). Ventana DISCOVERY inhibitor (8mins), casesin (1:10, 8mins) were then placed on the slides. With the slides warmed up to 37°C, the Cytochrome P450 2E1 antibody (1:100, Abcam, Ab28146) (60mins) and casesin (1:10, 1x 8mins) were used. Next, the slides were exposed to anti-rabbit IgG H (Roche DISCOVERY UltraMap, 760-4315) (10mins). To initiate the signal, the HRP-activated chromagen, DISCOVERY purple was added (40mins). Finally, Haemotoxylin, followed by bluing reagent was used as the counterstain.

#### Western blot

CYP2E1 and Dicer protein expression was quantified by western blot analysis. Protein homogenate was denatured at 70°C for 15 minutes and each sample was added at 30 µg to either 4-12% or 4-20 % tris-glycine precast gels (Novex™, Wedgewell™, Invitrogen). Following 1 hour of blocking with 5% milk (diluted in TSB-T wash buffer), the primary antibodies (**Table 1**) were added to 1% milk (dissolved in TBS-T) and incubated with the membranes (overnight, 4°C). The membranes were washed with TBS-T (3×10mins) and then transferred into the HRP-secondary antibody (**Table 1)** (diluted in 1% milk, 1 hour, at room temperature). The membranes were washed as previously described and incubated with ECL reagent (Pierce™, ThermoFisher Scientific) (5 mins, at room temperature). The membranes were exposed to X-Ray film (CL-Xposure film, ThermoFisher Scientific), developed (Compact 4x, Xograph) and scanned. After the film development, the membranes were incubated in acid stripping buffer (2×30 mins) and washed with PBS (3×3mins) ready to re-probe for β-actin following the methods above.

**Table 1.** Antibodies used in Western Blot experiments

| Target (immunogen) | Host and Conjugate | Source and Catalogue no. | Dilution factor |
| --- | --- | --- | --- |
| **Primary antibodies** |  |  |  |
| Dicer | Rabbit | Sigma Aldrich, SAB004200087 | 1:200 |
| Cytochrome P450 2E1 | Rabbit | Abcam, ab28146 | 1:5000 |
| β-actin | Mouse | Sigma Aldrich, A2228 | 1:20,000 |
| **Secondary antibodies** |  |  |  |
| Rabbit IgG | Goat, HRP | Vector Laboratories, PI-1000-1 | 1:3000 |
| Mouse IgG | Horse, HRP | Vector Laboratories, PI-2000-1 | 1:3000 |

#### In situ hybridisation

miRNA-122 was localised in the liver using the Discovery ULTRA Staining Module (Roche Diagnostics). During the staining programme, the miRCURY LNA miRNA Detection probes (mmu-miR-122-5p, YD00615338) and controls (U6, YD00699002 and Scramble-miR, YD00699004) were added. These were prepared using the miRCURY LNA miRNA ISH Buffer Set (FFPE) (Qiagen) mixed 1:1 with UltraPure DEPC water (Thermo Fisher Scientific).

#### Cytochrome p450 2E1 activity measurements

Tissue was homogenised into a microsomal preparation. The liver and kidney protein was diluted to 1mg/ml and 3mg/ml, respectively in phosphate buffer (pH 7.4). To determine the CYP2E1 metabolism activity, the sample was added to the CYP2E1 substrate, chlorzoxazone and NADPH mixture. The reaction was halted by a quench solution containing 200nM of benzoxazol and 4nM of verapamil 10 minutes after the incubation. Analyte measurements were made on the Xevo Water TQ-S Mass Spectrometry machine using the Water’s Masslynx software (Waters Corporation, Massachusetts, United States). Response of the CYP2E1 enzyme was determined from: 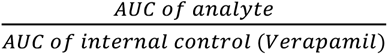

### Statistical Analysis

All data are presented as median±IQR. When two groups were compared the Mann-Whitney Test was used. For time course data on separate mice per time point linear regression was performed then each treatment was compared to null treatment to determine whether slopes were significantly different.

### Study approval

All animal studies were performed in accordance with the Animals (Scientific Procedures) Act 1986 / ASPA Amendment Regulations 2012 following ethical review by the University of Edinburgh. The local research ethics committee prospectively approved the human study, and informed consent was obtained from all patients before entry into the study.

## Author contributions

Experiments were performed by OM, EM, JT and PSL. Technical support was provided by IT, AS, RS, HR, KM. Supervision was performed by GAG, CG, KP, LD, ND, MB NH and DW. The work was directed by JD.

## Acknowledgments

Kidney Research UK Project Grant (RP_016_20170302) and Chest, Heart Stroke Scotland (CHSS) Project Grant (R15/A160) funded the work. CHSS funded the human studies. Author OM was supported by a Medical Research Scotland PhD studentship (Ref 875-2015). The MRC Scottish Clinical Pharmacology and Pathology Training Programme supported EM. JT was supported by a PhD studentship from the Cunningham Trust. JWD was supported by an NHS Research Scotland (NRS) Career Research Fellowship through NHS Lothian and acknowledges the contribution of the British Heart Foundation Centre of Research Excellence Award.

---

**Supplementary Figure 1.**
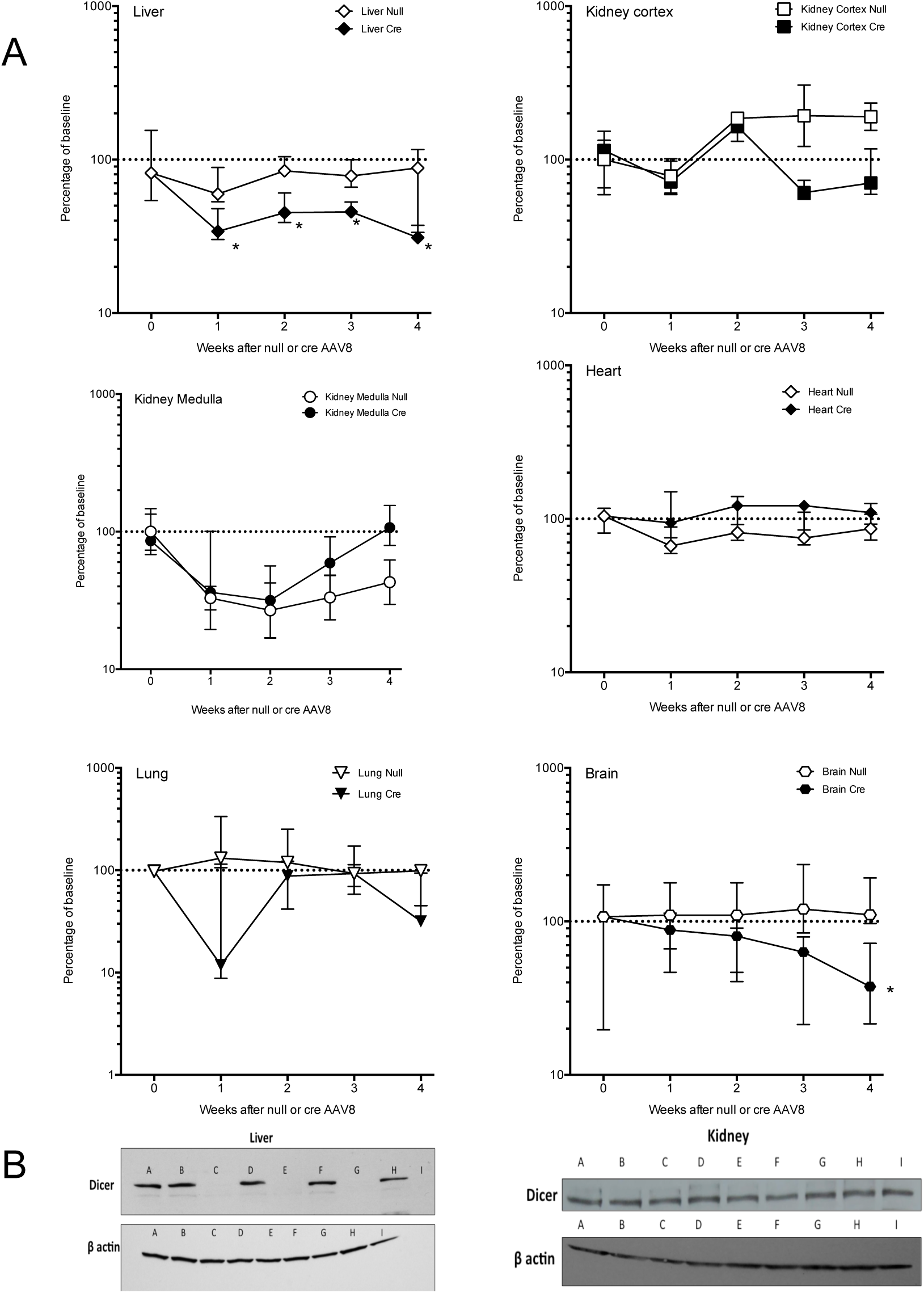
A The fold change over time in organ Dicer mRNA expression after treatment of Dicer^flox/flox^ mice with AAV8 vector expressing or not expressing Cre Recombinase (Cre). For each organ the expression of Dicer is expressed as a percentage of the untreated Dicer^flox/flox^ mice (baseline). Data are normalised to 18S. Filled symbols represent mice receiving Cre-AAV8. Unfilled symbols represent Null-AAV8. Symbols represent the time point median and the error bars define the inter-quartile range. N=20 for each AVV8/microRNA combination (N=5 for each time point). * = P<0.05 compared with baseline by Mann-Whitney Test. B Western blot of Dicer protein expression in liver and kidney. A = baseline untreated mouse. B, D, F, H AAV8 null treated mice and tissue collected 1, 2, 3 and 4 weeks later. C, E, G, I AAV8 Cre treated mice and tissue collected 1, 2, 3 and 4 weeks later.

**Supplementary Figure 2.**
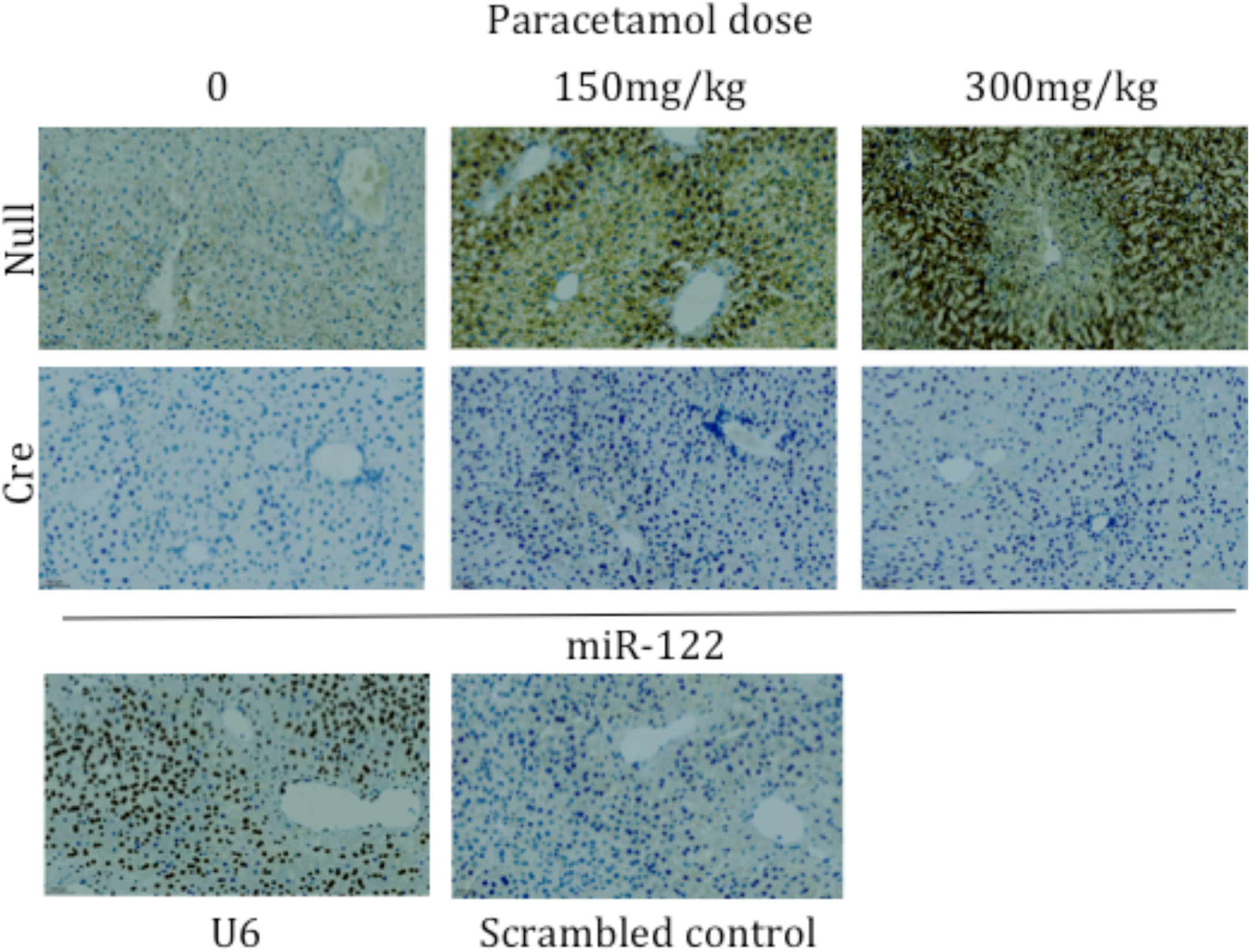
Liver injury due to paracetamol induces a change in miR-122 expression which is absent following conditional knock out of Dicer. DICER ^flox/flox^ mice were treated with AAV8 vector expressing or not expressing Cre Recombinase (Cre or Null). 3 weeks after AAV8 treatment mice received paracetamol 150 or 300 mg/kg (or vehicle (0)). In situ hybridisation for miR-122 in the liver was performed as described in methods. U6 and scrambled microRNA probe controls are also presented.

**Supplementary Figure 3.**
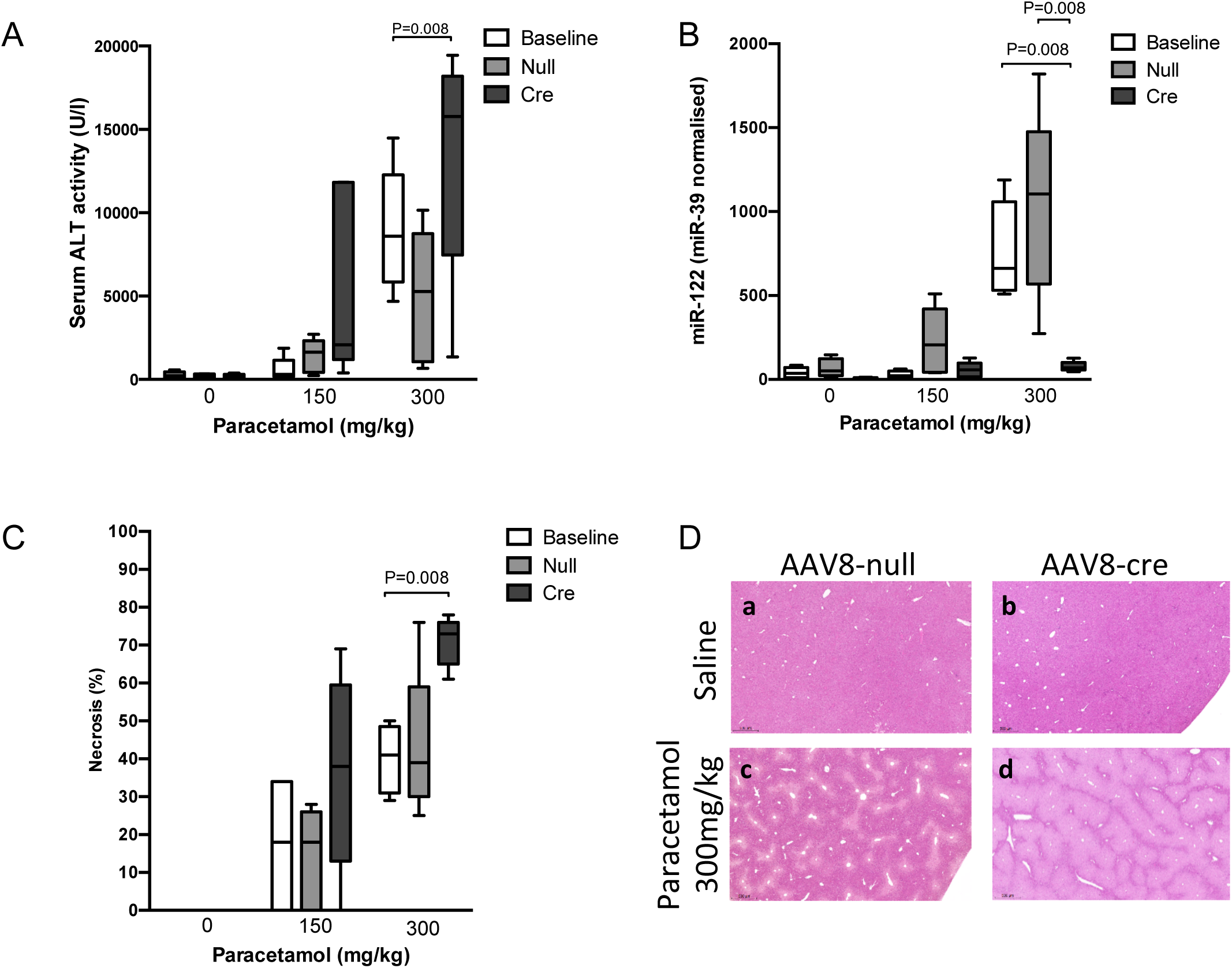
Dicer^flox/flox^ mice were treated with AAV8 vector expressing or not expressing Cre Recombinase (Cre or null). 3 weeks after AAV8 treatment mice received paracetamol 150 or 300 mg/kg (or vehicle (0)), then liver, kidney cortex and kidney medulla were harvested 6 hours later. Baseline = Dicer^flox/flox^ mice not receiving AAV8. Serum alanine transaminase activity (ALT) (A), serum miR-122 normalised to spike-in *C.elegans* miR-39 (B) and liver necrosis scores are presented in graphs (C). Liver necrosis scored as per methods. N=5 per group. Statistical significance was determined by Mann-Whitney Test. Data are represented as Tukey plots. Representative liver histology from AAV8-null and Cre treated Dicer^flox/flox^ mice are presented with vehicle or paracetamol treatment (D).

**Supplementary Figure 4.**
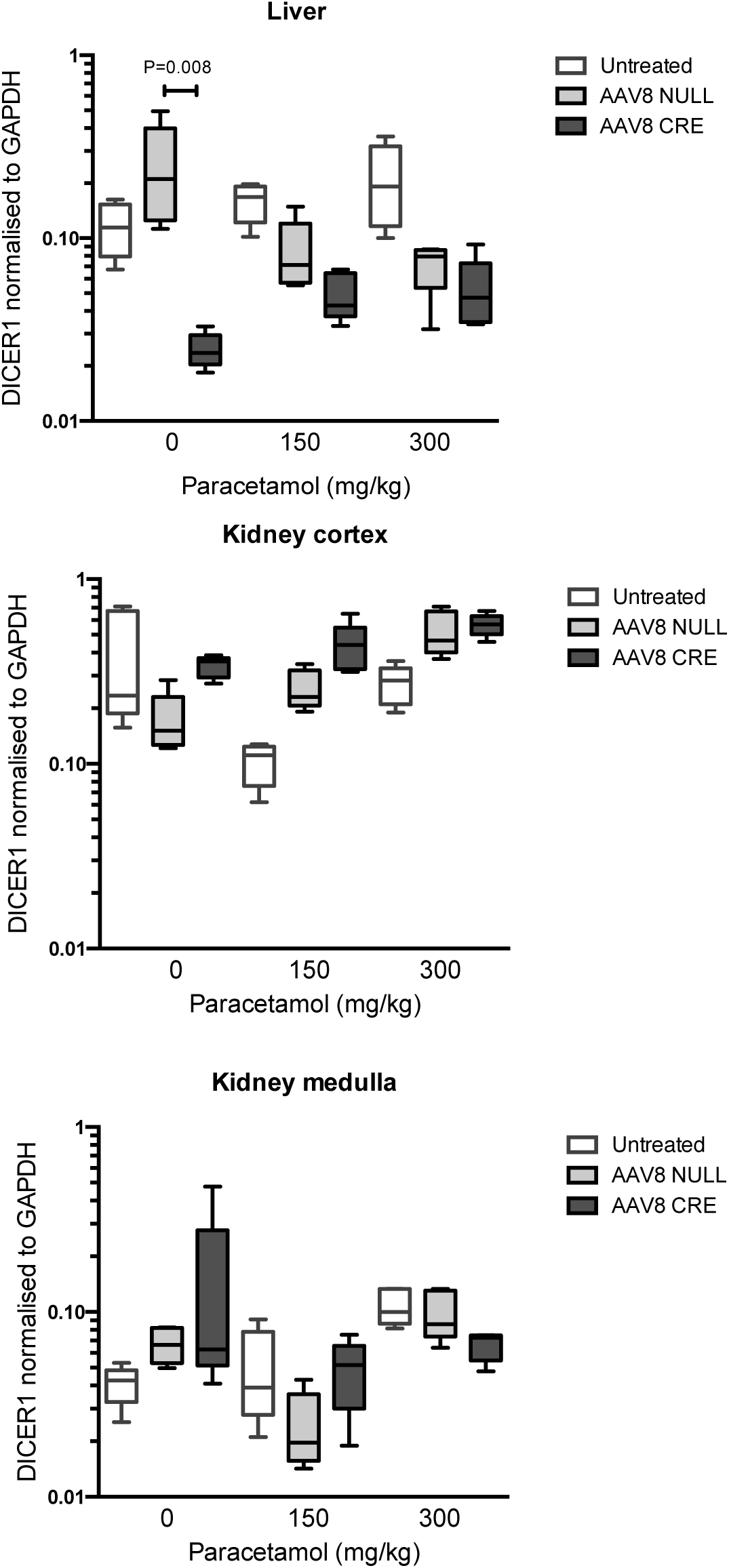
Dicer^flox/flox^ mice were treated with AAV8 vector expressing or not expressing Cre Recombinase (Cre or Null). 3 weeks after AAV8 treatment mice received paracetamol 150 or 300 mg/kg (or vehicle (0)) ip, then liver, kidney cortex and kidney medulla were harvested 6 hours later. Untreated = Dicer^flox/flox^ mice not receiving AAV8. Dicer mRNA expression is expressed as GAPDH(ct) - DICER (ct). N=5 per group. Statistical significance was determined by Mann-Whitney Test. Data are represented as Tukey plots.

**Supplementary Figure 5.**
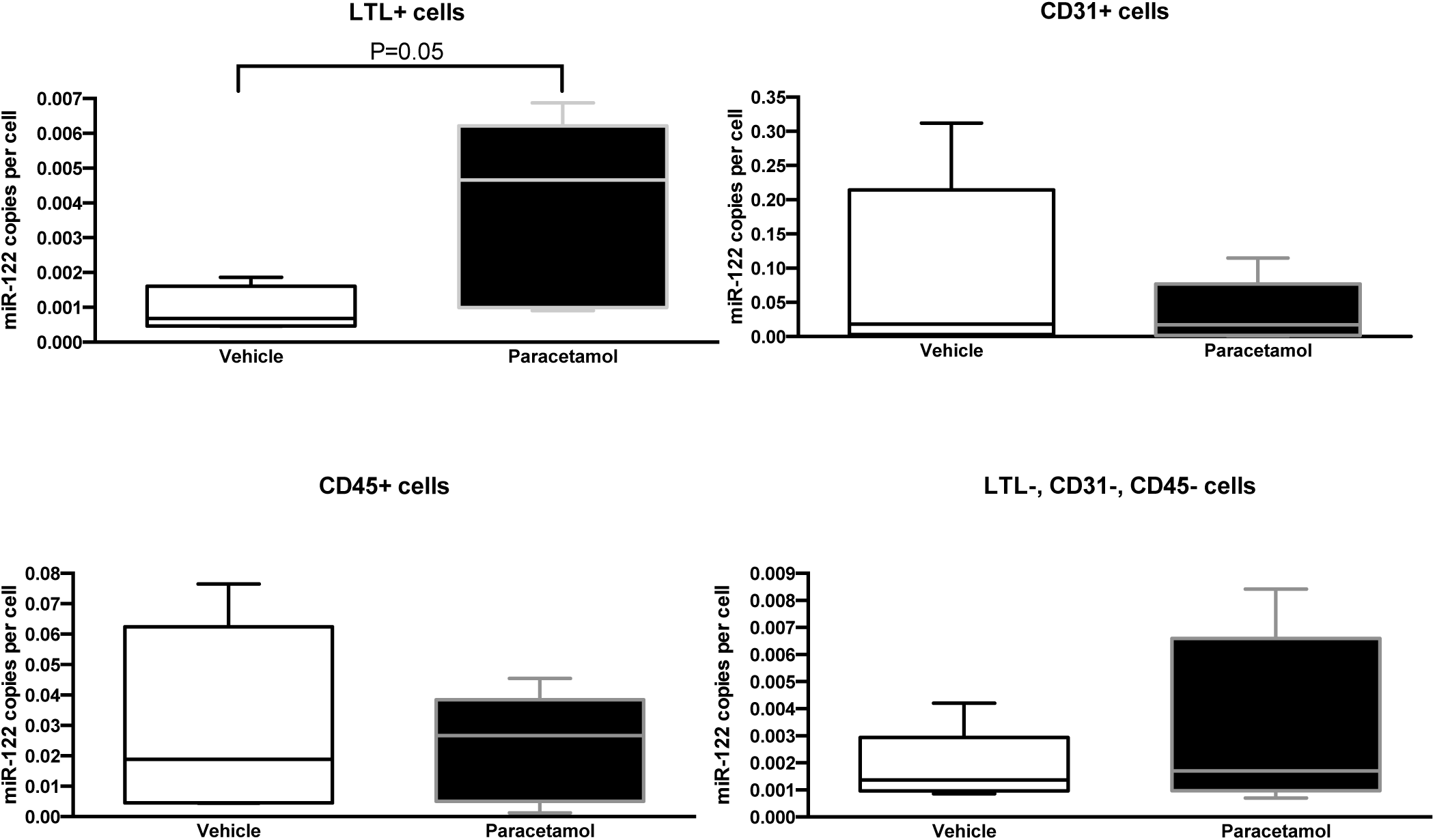
Six hours after treatment with paracetamol (300mg/kg ip) kidney cells were FACS sorted and miR-122 was measured by PCR. A standard curve was used to calculate the absolute copy number per cell. Statistical significance was determined by Mann-Whitney Test. Data are represented as Tukey plots.

**Supplementary Figure 6.**
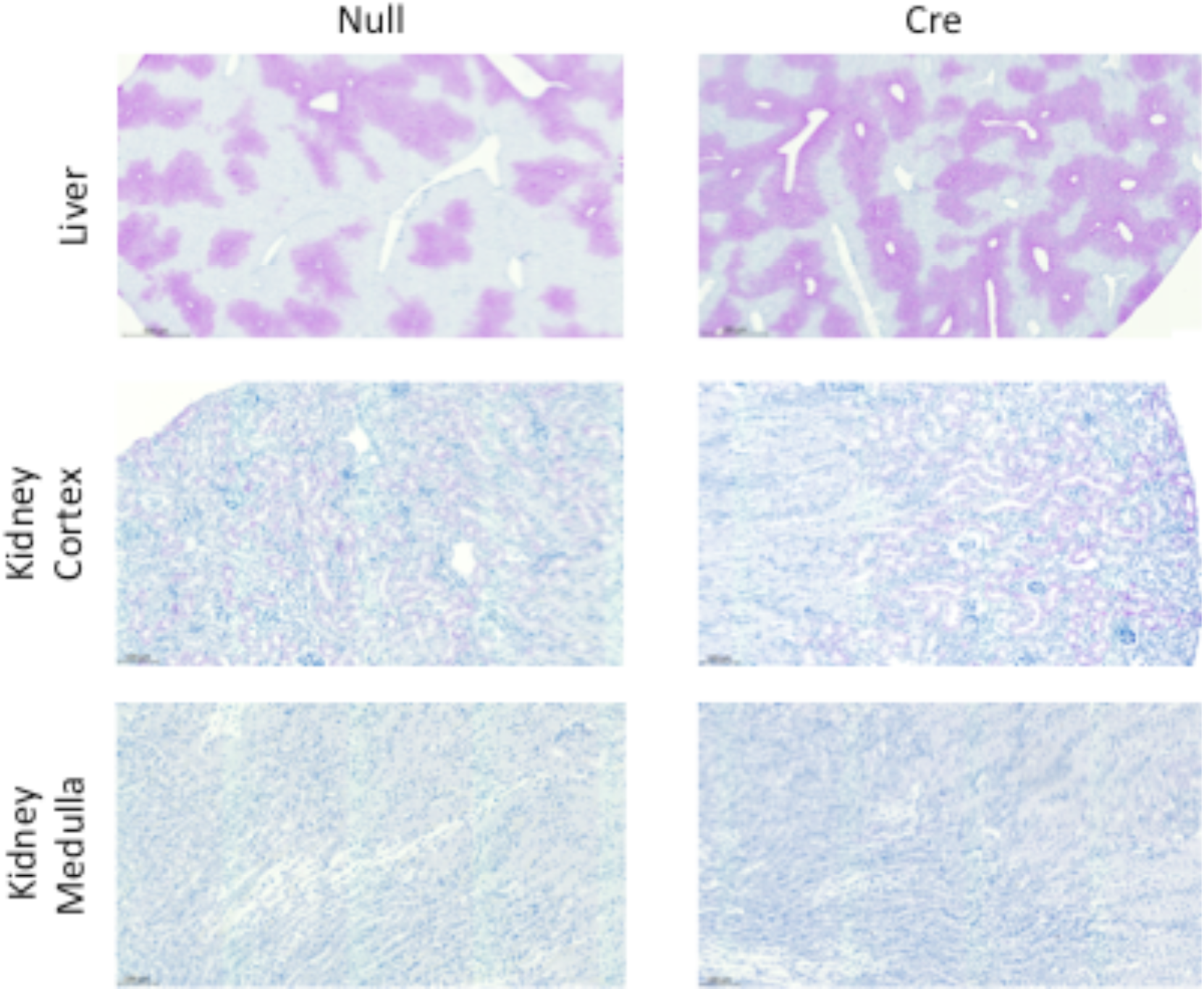
CYP2E1 is expressed in the liver and kidney cortex. DICER ^flox/flox^ mice were treated with AAV8 vector expressing or not expressing Cre Recombinase (Cre or Null). 3 weeks after AAV8 treatment mice tissue was collected for immunohistochemisty of CYP2E1.

## References

1. Bartel DP. Metazoan MicroRNAs. Cell 2018; 173(1): 20–51.

2. Mitchell PS, Parkin RK, Kroh EM, et al. Circulating microRNAs as stable blood-based markers for cancer detection. Proceedings of the National Academy of Sciences of the United States of America 2008; 105(30): 10513–8.

3. Wang K, Zhang S, Marzolf B, et al. Circulating microRNAs, potential biomarkers for drug-induced liver injury. Proceedings of the National Academy of Sciences of the United States of America 2009; 106(11): 4402–7.

4. Valadi H, Ekstrom K, Bossios A, Sjostrand M, Lee JJ, Lotvall JO. Exosome-mediated transfer of mRNAs and microRNAs is a novel mechanism of genetic exchange between cells. Nature cell biology 2007; 9(6): 654–9.

5. Fong MY, Zhou W, Liu L, et al. Breast-cancer-secreted miR-122 reprograms glucose metabolism in premetastatic niche to promote metastasis. Nature cell biology 2015; 17(2): 183–94.

6. Oosthuyzen W, Scullion K, Ivy JR, et al. Vasopressin regulates extracellular vesicle uptake by kidney collecting duct cells. J Am Soc Nephrol 2016; 27(11): 3345–3355.

7. Pakravan N, Simpson K, Waring WS, Bates CM, Bateman DN. Renal injury at first presentation as a predictor for poor outcome in severe paracetamol poisoning referred to a liver transplant unit. Eur J Clin Pharmacol 2009; 65(2): 163–8.

8. Jopling C. Liver-specific microRNA-122: Biogenesis and function. RNA Biol 2012; 9(2): 137–42.

9. de Rie D, Abugessaisa I, Alam T, et al. An integrated expression atlas of miRNAs and their promoters in human and mouse. Nat Biotechnol 2017; 35(9): 872–8.

10. Starkey Lewis PJ, Dear J, Platt V, et al. Circulating microRNAs as potential markers of human drug-induced liver injury. Hepatology 2011; 54(5): 1767–76.

11. Dear JW, Clarke JI, Francis B, et al. Risk stratification after paracetamol overdose using mechanistic biomarkers: results from two prospective cohort studies. Lancet Gastroenterol Hepatol 2018; 3(2): 104–13.

12. Jaeschke H, Xie Y, McGill MR. Acetaminophen-induced Liver Injury: from Animal Models to Humans. J Clin Transl Hepatol 2014; 2(3): 153–61.

13. Gill P, Bhattacharyya S, McCullough S, et al. MicroRNA regulation of CYP 1A2, CYP3A4 and CYP2E1 expression in acetaminophen toxicity. Sci Rep 2017; 7(1): 12331.

14. Chowdhary V, Teng KY, Thakral S, et al. miRNA-122 Protects Mice and Human Hepatocytes from Acetaminophen Toxicity by Regulating Cytochrome P450 Family 1 Subfamily A Member 2 and Family 2 Subfamily E Member 1 Expression. Am J Pathol 2017; 187(12): 2758–74.

15. Jones AF, Vale JA. Paracetamol poisoning and the kidney. J Clin Pharm Ther 1993; 18(1): 5–8.

16. Liu H, Baliga R. Cytochrome P450 2E1 null mice provide novel protection against cisplatin-induced nephrotoxicity and apoptosis. Kidney international 2003; 63(5): 1687–96.

17. Vliegenthart AD, Shaffer JM, Clarke JI, et al. Comprehensive microRNA profiling in acetaminophen toxicity identifies novel circulating biomarkers for human liver and kidney injury. Sci Rep 2015; 5: 15501.

18. Bala S, Petrasek J, Mundkur S, et al. Circulating microRNAs in exosomes indicate hepatocyte injury and inflammation in alcoholic, drug-induced, and inflammatory liver diseases. Hepatology 2012; 56(5): 1946–57.

19. Rivkin M, Simerzin A, Zorde-Khvalevsky E, et al. Inflammation-Induced Expression and Secretion of MicroRNA 122 Leads to Reduced Blood Levels of Kidney-derived Erythropoietin and Anemia. Gastroenterology 2016; 151(5): 999–1010.

20. Wang Y, Liang H, Jin F, et al. Injured liver-released miRNA-122 elicits acute pulmonary inflammation via activating alveolar macrophage TLR7 signaling pathway. Proceedings of the National Academy of Sciences of the United States of America 2019; 116(13): 6162–71.

21. John K, Hadem J, Krech T, et al. MicroRNAs play a role in spontaneous recovery from acute liver failure. Hepatology 2014; 60(4): 1346–55.

22. Arroyo JD, Chevillet JR, Kroh EM, et al. Argonaute2 complexes carry a population of circulating microRNAs independent of vesicles in human plasma. Proc Natl Acad Sci U S A 2011; 108(12): 5003–8.

23. Oosthuyzen W, Sime NE, Ivy JR, et al. Quantification of human urinary exosomes by nanoparticle tracking analysis. J Physiol 2013; 591(Pt 23): 5833–42.

24. Michael LH, Entman ML, Hartley CJ, et al. Myocardial ischemia and reperfusion: a murine model. Am J Physiol 1995; 269(6 Pt 2): H2147–54.

25. Looi YH, Grieve DJ, Siva A, et al. Involvement of Nox2 NADPH oxidase in adverse cardiac remodeling after myocardial infarction. Hypertension 2008; 51(2): 319–25.

